# p97/VCP targets *Toxoplasma gondii* vacuoles for parasite restriction in interferon-stimulated human cells

**DOI:** 10.1101/2023.06.20.545566

**Authors:** Barbara Clough, Daniel Fisch, Todd H Mize, Vesela Encheva, Ambrosius Snijders, Eva-Maria Frickel

## Abstract

Infection with the parasite *Toxoplasma gondii* leads to production of interferon gamma (IFNγ) that stimulates cells to upregulate defence proteins targeting the parasite for cell intrinsic elimination or growth restriction. Various host defence mechanisms operate at the parasitophorous vacuole (PV) in different human cell types leading to PV disruption, acidification, or membrane envelopment. Ubiquitin and p62 are players in all human host control mechanisms of *Toxoplasma*, but other unifying proteins have not been identified. Here, we show that p97/valosin-containing protein (VCP), as well as its associated proteins ANKRD13A and UBXD1 control *Toxoplasma* infection while recruited to the PV in IFNγ-stimulated endothelial cells. Convergent deposition of ANKRD13A, p97/VCP and UBXD1 onto the same vacuole is dependent on vacuolar ubiquitination and observed within 2h post-infection. ANKRD13A, p97/VCP and UBXD1 all drive the acidification mechanism of the vacuole, which is the IFNγ-dependent control pathway of *Toxoplasma* in endothelial cells. We assessed p97/VCP in *Toxoplasma* control in various human cells and demonstrate that p97/VCP is a universal IFNγ-dependent host restriction factor targeting the *Toxoplasma* PV in epithelial (HeLa) and endothelial cells (HUVEC), fibroblasts (HFF) and macrophages (THP1).

## 1. INTRODUCTION

*Toxoplasma gondii* (Tg) is a global human pathogen infecting one third of the world’s population (Pappas et al., 2009). Parasite infection leads to the production of the inflammatory cytokine interferon gamma (IFNγ) by natural killer (NK) and T cells. IFNγ acts on host cells to upregulate defence proteins that control the parasite and, in some cases, induce host cell death to eliminate the host replicative niche. Key to the success of Tg is its ability to replicate within many host species and infect most nucleated cells (Jones et al., 1972). The intracellular lifestyle of Tg is further protected as the parasite forms a vacuolar compartment from the host plasma membrane upon invasion, the parasitophorous vacuole (PV), which it uses as a strong-hold to mount counterattacks against its host (Clough and Frickel, 2017).

IFNγ has long been shown to be the major player in the control of Tg with studies in mice revealing a dose-dependent effect of monoclonal antibodies to IFNγ on survival after Tg infection (Suzuki et al., 1988). The control of Tg tachyzoites in mice operates via mechanisms common to all murine cell types, involving two families of IFNγ-dependent GTPases, the immunity-related GTPases (IRGs) and guanylate binding proteins (GBPs), with some Tg strains able to resist IRG-driven elimination. These GTPase families accumulate and co-ordinate a sequential, IRG-led attack on the PV resulting in vacuolar breakage and parasite death (Khaminets et al., 2010). In contrast to murine infection, study of Tg infection in human cells has revealed a complex interplay between host defence and Tg attack, with different cell types employing distinct mechanisms to combat the pathogen (Saeij and Frickel, 2017, Frickel and Hunter, 2021). Many of the control mechanisms are dependent on the Tg strain-and IFNγ-dependent recruitment of ubiquitin to the PV within minutes of Tg invasion (Clough et al., 2016, Selleck et al., 2015). However, distinct from murine Tg infection, all Tg strains studied to date are eliminated in human cells.

Our study centres on understanding the cell type specific differences in Tg elimination in ubiquitin-marked vacuoles. Following ubiquitin targeting, host defence diverges to include recruitment of further host proteins to type II and III parasite vacuoles, including the autophagy-associated and ubiquitin-binding proteins p62 and NDP52. Host E3 ligases TRAF2 and TRAF6 have been implicated in bringing p62, NDP52, LC3B and GABARAP to the Tg PV in IFNγ-stimulated primary HFFs, mediated by the parasite secreted protein GRA15 (Mukhopadhyay et al., 2020). In the HeLa epithelial cell line, LC3 and GABARAPL2 also undergo ubiquitin-dependent accumulation at the PV leading to Tg growth restriction (Selleck et al., 2015, Zhang et al., 2020). This contrasts with primary HUVEC endothelial cells where vacuoles become LAMP1 positive and acidify leading to parasite death via a non-autophagic mechanism (Clough et al., 2016). In the latter two scenarios, the PV does not to break, distinguishing these mechanisms strikingly from the IFNγ-driven murine host cell defence. In human cells vacuole breakage has only been observed in immune cells such as the THP-1 macrophage cell line and primary blood monocyte-derived macrophages. Here, the vacuole is also ubiquitinated and subsequently decorated with GBP1 in type I and II strains of Tg, resulting in breakage of the PV membrane, culminating in cell death through an apoptotic pathway (Fisch et al, unpublished and (Fisch et al., 2020)). More recently, in both A549 and HFF cells, the IFNγ-mediated control of *Toxoplasma* has been shown to be directed primarily by the IFNγ-inducible host E3-ligase RNF213, which promotes Lys63 and M1 ubiquitination at the PV leading to the recruitment of host defence proteins p62, NDP52, optineurin and TAX1BP1 (Hernandez et al., 2022).

Host defence mechanisms also operate distal to the PV in human cell lines. Prior research has shown that IFNγ-dependent control of Tg in human fibroblasts could be attributed to tryptophan starvation of the tryptophan-auxotrophic Tg, through induction of host indoleamine 2,3-dioxygenase (IDO1) (Pfefferkorn, 1984, Pfefferkorn et al., 1986). Classically viewed as PV-targeting host defence proteins, GBPs have been shown to also operate in PV-distal control of Tg. GBP1 is not observed at the vacuolar membrane in the lung epithelial cell line A549 but is instrumental in controlling parasite survival (Johnston et al., 2016). GBP2 and GBP5 do not target the PV in human macrophages yet are able to restrict parasite growth in a GTPase-dependent fashion (Fisch et al., 2021a). This suggests a more complex interplay of host defence mechanisms working both distal to and at the vacuolar membrane.

The cell-specific molecular mechanisms of parasite control will become clearer as more host proteins are identified which co-operate with these ubiquitin-dependent host factors. Here, we identify p97/ valosin-containing protein (VCP) as a protein that targets the Tg vacuole and operates in host defence in a range of human cell types.

## 2. RESULTS

### 2.1 ANKRD13A is a ubiquitin substrate in *Toxoplasma*-infected human cells

The PV of Tg type II Pru in epithelial and endothelial cells is ubiquitinated in a parasite strain-and IFNγ-dependent manner (Figure S1a and (Clough et al., 2016, Selleck et al., 2015)), with Lys63-linked ubiquitin being one of the dominant linkage types observed (Clough et al., 2016, Hernandez et al., 2022). Ubiquitination is a key step in the recognition and tagging of parasite vacuoles for destruction, with the linkage pattern of ubiquitination determining the fate and function of the ubiquitinated substrates (Ikeda and Dikic, 2008).

We took a proteomics approach using stable isotope labelling by amino acids in cell culture (SILAC) with mass spectrometry to identify new candidate ubiquitinated host proteins at the PV (Figure S1b). Comparison of IFNγ-stimulated A549 epithelial cells, Tg-infected versus uninfected cells, was made using SILAC. Enrichment of ubiquitinated peptides by di-glycine immunoprecipitation was performed after tryptic digestion to enable quantitation of the changes in ubiquitination of host and parasite components on infection. This was followed by proteomic analysis using liquid chromatography coupled to tandem mass spectrometry (LC-MS/MS). A list of ubiquitinated host peptides specific for cells infected with Tg type II Pru was compiled with targets likely to be present at the PV identified from a distribution plot of target proteins (Figure S1c and Table S1). We show that a host protein, ANKRD13A, is ubiquitinated on host cell infection by Tg type II Pru (Figure S1c). ANKRD13A was of particular interest since this protein had been shown specifically to bind Lys63-linked ubiquitin chains and to be involved in membrane protein modulation and trafficking mechanisms (Tanno et al., 2012, Burana et al., 2016).

### 2.2 ANKRD13A, p97/VCP and UBXD1 control *Toxoplasma* localised to the parasitophorous vacuole

ANKRD13A has been shown to interact with the AAA+ ATPase p97/VCP, which has many and diverse cellular roles with ubiquitinated substrate proteins being a common target for its functions (Stach and Freemont, 2017, Burana et al., 2016). Consequently, we wanted to address the fate of Tg in cells depleted of ANKRD13A and p97/VCP and the p97/VCP co-factor UBXD1. A combination of siRNAs for each of the proteins were transfected into HUVEC by nucleofection along with a non-targeting control siRNA. A marked reduction in all proteins was seen with the siRNA treatment by immunoblotting compared to the control siRNA (Figure S2a). The percentage of infected cells and vacuole to cell ratio of Tg type II Pru were counted at 6h post infection by automated high content imaging (HCI) followed by host response to microbe analysis (HRMAn) (Figure 1a; (Fisch et al., 2021b)). A significant increase in both the percentage of infected cells and vacuole to cell ratio was observed in cells where ANKRD13A, p97/VCP or UBXD1 was knocked down (Figure 1a). Furthermore, recruitment of all target proteins to Tg vacuoles was observed at 2h p.i. for Tg type II Pru, with a significant increase in recruited vacuoles after stimulation of HUVEC with IFNγ (Figure 1b). Similar observations were made for Tg type III CEP (Figure 1c and d). To observe the kinetics of p97/VCP recruitment, live imaging was performed 2-3h post-infection of EGFP-p97/VCP-transfected, IFNγ-stimulated HUVEC by Td-tomato stably transfected Tg type II Pru (Figure S3a and Video S1). p97/VCP was observed to accumulate at the vacuole, and in one sequence the parasite within the targeted PV loses fluorescence indicating a loss of parasite viability and possible vacuolar acidification (Figure S3ai and Video S1).

**Figure 1.**
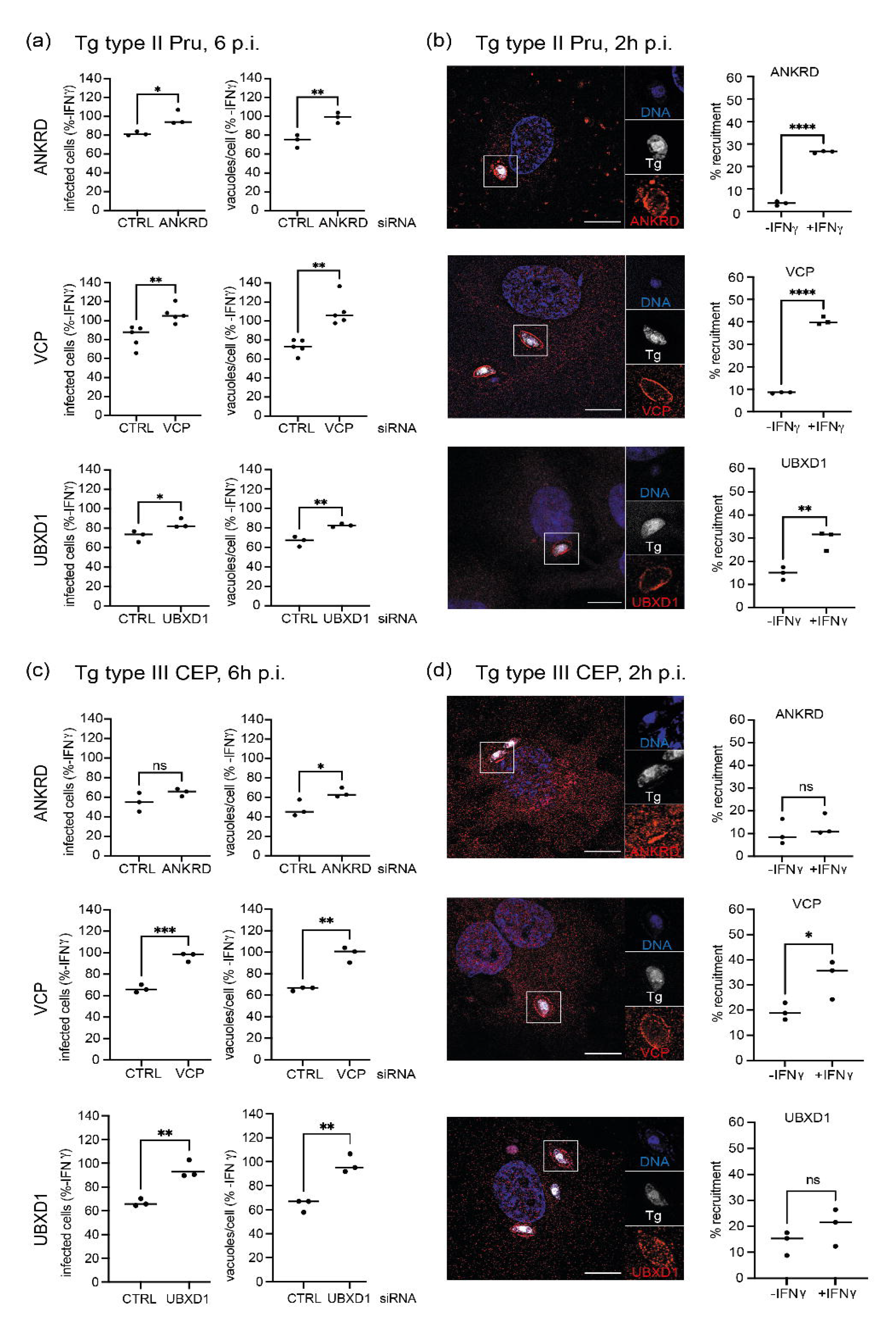
ANKRD13A, p97/VCP and UBXD1 target *Toxoplasma* (Tg) for interferon gamma (IFNγ)-driven restriction. **a)** Target gene knockdown leads to an increased percentage of infected cells and more vacuoles/cell in IFNγ-stimulated HUVEC 6h p.i. with Tg type II Pru compared to unstimulated cells. >1000 cells counted. n = 3-5. **b)** Representative structured illumination microscopy (SIM) image and quantification of target protein (red) recruitment to Tg type II Pru (white) vacuoles in HUVEC in dependency of IFNγ stimulation 2h p.i. >200 vacuoles counted. Scale bar = 10μm. **c)** Target gene knockdown leads to an increased percentage of infected cells and more vacuoles/cell in IFNγ-stimulated HUVEC 6h p.i. with Tg type III CEP compared to unstimulated cells. >1000 cells counted. n = 3. **d)** Representative SIM image and quantification of target protein (red) recruitment to Tg type III CEP (white) vacuoles in HUVEC in dependency of IFNγ stimulation 2h p.i. Scale bar =10μm.

We conclude from these data that the host proteins ANKRD13A, VCP and UBXD1 are important in controlling the survival of Tg type II Pru and type III CEP and mediate their effect at the Tg vacuole.

### 2.3 ANKRD13A, p97/VCP and UBXD1 are ubiquitin-dependent host factors that target a PV subset

Next, we wanted to determine whether ubiquitin was the driving factor for recruitment of ANKRD13A, VCP and UBXD1 and whether these host proteins were directed to the same subset of PV. Recruitment of total ubiquitin, p97/VCP, ANKRD13A and UBXD1 at 2h was determined at the PV in cells pre-stimulated with IFNγ and then treated for 2h with the E1 enzyme inhibitor, PYR41, prior to infection with Tg type II Pru. Without PYR41, IFNγ drives the deposition of ubiquitin onto the PV, and a reduction in ubiquitin recruitment in HUVEC stimulated with IFNγ confirmed the activity of the inhibitor (Fig 2a). We observed significant decreases in recruitment of VCP, ANKRD13A and UBXD1 in IFNγ-stimulated cells, revealing that ubiquitin was necessary for PV targeting by these host factors (Fig 2b). ANKRD13A has been shown to bind Lys63-ubiquitinated substrates via ubiquitin-interaction motifs (UIMs) 3 and 4 located at its C-terminal end, with the interaction being regulated by its own monoubiquitylation (Tanno et al., 2012). Using cells transfected with WT-ANKRD13A and mutants deleted in the C-terminal UIM domain, we found that mutants lacking UIMs 3 and 4 did not localise at all to Lys63-ubiquitin positive vacuoles (Figure 2c). Furthermore, a lack of PV targeting was observed for the ANKRD13A UIM1-4 deletion mutant (Figure 2c). Deletion of ANKRD13A’s UIM domains prevented observable accumulation of Lys63-ubiquitin at the PV (Figure 2c).

**Figure 2.**
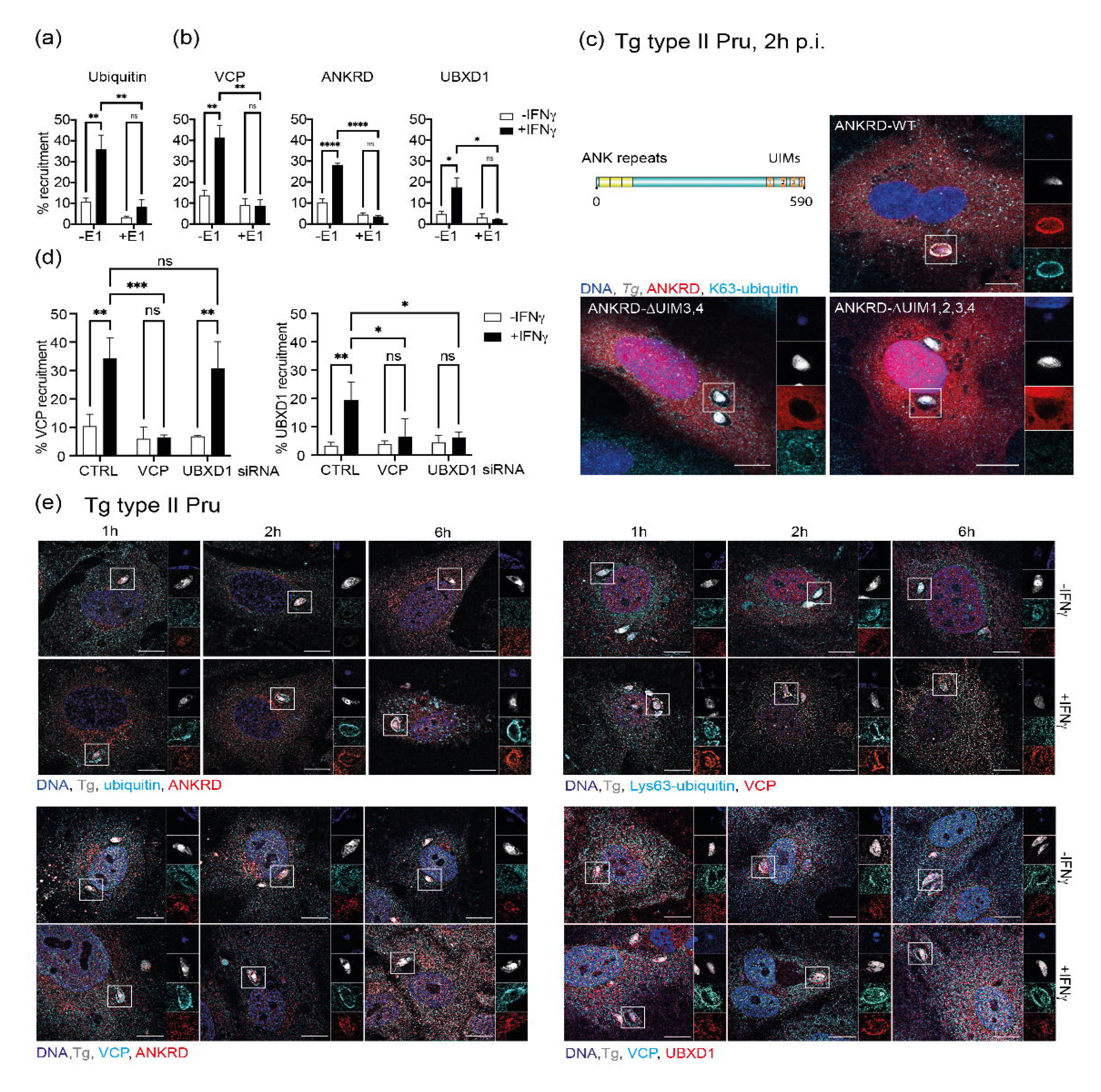
Ubiquitin-dependent recruitment of ANKRD13A, p97/VCP and UBXD1 colocalises to a subset of *Toxoplasma* (Tg) vacuoles. **a)** Total ubiquitin, VCP, ANKRD13A and UBXD1 are recruited to Tg type II Pru vacuoles in HUVEC 2h p.i., in dependency of interferon gamma (IFNγ) stimulation. Treatment of cells with E1 ubiquitination inhibitor PYR41 (+E1) significantly reduces the recruitment of these target proteins to Tg type II Pru vacuoles. >200 vacuoles counted, n=3. **b)** ANKRD13A is recruited to Tg type II Pru vacuoles through its ubiquitin interaction motif (UIM) domains. Domain structure of ANKRD13A (image created in BioRender.com) showing UIM domains with representative confocal images depicting over-expression of ANKRD13A UIM deletion mutants (red) and anti-Lys63- ubiquitin staining (cyan). Scale bar = 10μm. **c)** UBXD1 accumulation at Tg type II Pru vacuoles 2h p.i. in IFNγ-stimulated HUVEC is dependent on p97/VCP recruitment, while VCP is able to target vacuoles in the absence of UBXD1. Recruitment of p97/VCP and UBXD1 to >200 Tg type II Pru vacuoles was determined at 2h p.i. in cells knocked down for either p97/VCP or UBXD1. n=3. **d)** Host proteins target the same subset of Tg type II Pru vacuoles. Time course of denoted protein (red, cyan) co-recruitment to the Tg type II Pru (white) vacuole in HUVEC with or without IFNγ stimulation at 1, 2 and 6h p.i. Representative structured illumination microscopy (SIM) images are shown. Scale bar =10 μm.

As p97/VCP and UBXD1 are known to be co-factors for the endo-lysosomal damage response (ELDR) (Papadopoulos et al., 2017), we wanted to determine their respective hierarchy of recruitment to the PV. Knock downs of p97/VCP and UBXD1 were made in HUVEC using siRNA and recruitment of these proteins was scored at 2h p.i., with specific protein recruitment to corresponding siRNA knock down cells included as negative controls. While p97/VCP was still able to recruit to the PV in the absence of UBXD1 the converse was not observed (Figure 2d). This showed that UBXD1 recruitment was only possible in the presence of p97/VCP. Since UBXD1 targeting to the PV requires both p97/VCP and ubiquitin, it implies that the UBXD1 association with p97/VCP occurs subsequent to p97/VCP localisation to the PV. To assess whether these host proteins targeted the same PV, co-localisation studies were performed at 1, 2 and 6h post infection with Tg type II Pru (Figure 2e). Deposition of host proteins onto the PV was absent or limited in the absence of IFNγ (Figure 2e). This aligned with the slightly higher recruitment of these proteins in unstimulated cells observed in Figure 1b. Ubiquitin and p97/VCP recruitment to type II PV in IFNγ-treated cells was apparent from 1h p.i. with subsequent accumulation of ANKRD13A and UBXD1 from 2h, to ubiquitin and p97/VCP positive vacuoles (Figure 2e). These data reveal that ubiquitin, ANKRD13A, p97/VCP and UBXD1 are targeted to the same subset of vacuoles in IFNγ-stimulated cells with a hierarchy of recruitment. We show that ubiquitin drives recruitment to the vacuole during the first hour p.i. with p97/VCP accumulating on ubiquitinated PV from 1h p.i. and subsequent targeting of p97/VCP positive PVs by its co-factor UBXD1 and ANKRD13A after 2h p.i.

### 2.4 ANKRD13A, p97/VCP and UBXD1 drive PV acidification with p97/VCP being a universal *Toxoplasma* PV-localised IFN**γ**- dependent human host protein

Previously we reported that Tg type II Pru is targeted for destruction in IFNγ- stimulated HUVEC by endo-lysosomal acidification, initiated by ubiquitination at the PV (Clough et al., 2016). To address whether the new host factors ANKRD13A, p97/VCP and UBXD1 contributed to vacuolar acidification, we infected cells knocked down for each of the host proteins with Tg type II Pru for 6h, adding LysoTracker to the cultures 1h before fixation for microscopy. LysoTracker-stained vacuoles were clearly visible for parasite-infected control cells stimulated with IFNγ, whereas PV acidification was diminished in infected cells lacking the specified host proteins (Figure 3a, Figure S4). This supported the notion that ANKRD13A, p97/VCP and UBXD1 are involved in clearance of Tg by acidification of the PV.

**Figure 3.**
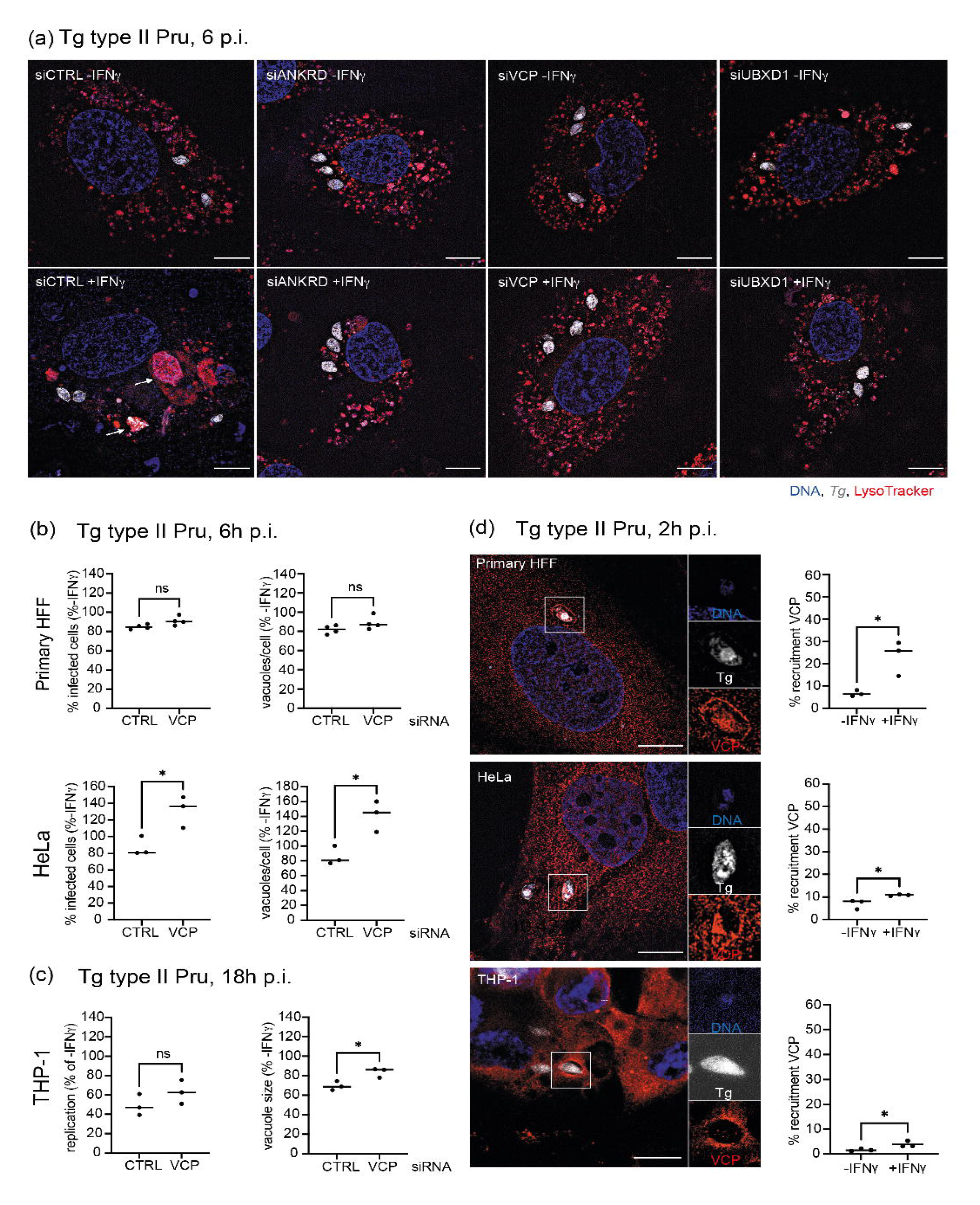
ANKRD13A, p97/VCP and UBXD1 are host defence proteins driving PV acidification in HUVEC with p97/VCP impacting *Toxoplasma* (Tg) in different cell types. **a)** Host proteins ANKRD13A, VCP and UBXD1 are involved in acidifying the Tg type II Pru vacuole in interferon gamma (IFNγ)-stimulated cells. Representative structured illumination microscopy (SIM) images of LysoTracker (red)-stained, Tg type II Pru (white) vacuoles 6h p.i. in HUVEC knocked down with control (siCTRL) or target gene siRNA with and without IFNγ. White arrows indicate examples of acidified vacuoles. Scale bar = 10μm. **b)** p97/VCP knockdown did not lead to a significant increase in percentage of infected cells or vacuoles/cell in IFNγ-stimulated primary fibroblasts (HFF) infected with Tg type II Pru at 6h p.i. compared to unstimulated cells. Depletion of p97/VCP led to an increased percentage of infected cells and more vacuoles/cell in IFNγ- stimulated human epithelial cells (HeLa) infected with Tg type II Pru at 6h p.i. compared to unstimulated cells. >1000 cells counted per condition, n=3. **c)** In the human macrophage (THP1) cell line, p97/VCP knockdown led to an increased vacuole size in IFNγ-stimulated cells with a trend towards increase replication. For THP1, readouts of % replication and vacuole size were determined at 18h p.i. since the level of cell death occurring in this cell type on infection interfered with analyses related to cell number. >1000 cells counted per condition, n=3. **d)** IFNγ-dependent recruitment of VCP/p97 to Tg type II Pru vacuoles was observed at 2h p.i. in primary HFF, HeLa and THP-1 cells. Representative SIM image are shown with quantitation. Scale bar = 10μm. >200 vacuoles were counted, n=3.

Since it has been shown that the mechanisms of parasite clearance vary in immune-stimulated human cells dependent upon cell type (Krishnamurthy et al., 2017), we were curious to see whether the host factor p97/VCP was an important defence molecule in different cell types. p97/VCP has been previously implicated in intracellular pathogen survival (Humphreys et al., 2009, Dorer et al., 2006) and is further involved in diverse cellular processes involving membrane protein recycling and degradation in a range of cell types (Stach and Freemont, 2017). Here, primary fibroblast (HFF) and endothelial (HeLa) cells knocked down for p97/VCP were infected with Tg type II Pru and the impact on percentage infected cells and vacuoles/cell determined at 6h p.i. using HRMAn analysis. While p97/VCP showed a significant effect on the IFNγ-dependent restriction of Tg in HeLa cells, there was little change observed in primary fibroblasts (Figure 3b). This indicated that at this time point p97/VCP was important for the IFNγ-dependent control of Tg type II Pru in HeLa, but not in primary HFFs. As THP-1 cells have been reported to undergo cell death upon IFNγ-induction and infection with Tg (Fisch et al., 2019), we analysed the infection readouts of % replication and vacuole size at 18h p.i.; parameters that do not depend upon total cell number. A significant increase in vacuole size was noted for immune-stimulated cells lacking p97/VCP, indicating that p97/VCP impacted the parasite’s ability to divide in macrophages (Figure 3c). We found that p97/VCP was recruited to Tg type II Pru vacuoles to different extents for primary fibroblasts, endothelial and macrophage cell lines (Figure 3d). IFNγ-dependent accumulation of p97/VCP increased significantly in all cell types with levels in primary fibroblasts higher than the other cell types.

Collectively these data show that ANKRD13A, p97/VCP and UBXD1 are host defence proteins that drive Tg type II Pru PVs to acidification in endothelial cells. p97/VCP is universally recruited to different human cell types in dependence of IFNγ, with diverse effects on Tg restriction.

#### Take Aways

- ANKRD13A, p97/VCP and UBXD1 are host factors that control Tg at the PV in endothelial cells.
- PV ubiquitination is a pre-requisite for recruitment of ANKRD13A, p97/VCP and UBXD1 in endothelial cells.
- PV deposition of these host defence proteins directs Tg PVs to acidification in endothelial cells.
- p97/VCP universally targets PVs in human cells and restricts Tg in different human cell types.

## 3. DISCUSSION

Ubiquitination of the PV is a universal priming step in the IFNγ-driven control of *Toxoplasma* in human cells (Clough and Frickel, 2017). We and others previously showed that ubiquitin-binding proteins p62 and NDP52 locate to the ubiquitinated PV and contribute to the elimination of Tg type II (Clough et al., 2016, Selleck et al., 2015). Here, we sought to examine pathways of Tg control that are seeded by PV ubiquitination. To this end, we employed mass spectrometry to identify ANKRD13A as a novel ubiquitinated protein during Tg infection in IFNγ-stimulated epithelial cells. We explored this further by studying the ANKRD13A-associated proteins p97/VCP and UBXD1 at the PV and show that they act in defence against Tg infection in IFNγ- stimulated human cells. We demonstrate that these host defence proteins collaborated in the vacuolar acidification of Tg in endothelial cells (Clough et al., 2016). Furthermore, p97/VCP was recruited to the Tg PV in a diverse range of human cell types with a differential effect on Tg control. This implies that although universally recruited to the Tg PV in endothelial, epithelial, macrophage and fibroblast cells, p97/VCP executes pathogen-defence functions in different human cell types possibly via divergent mechanisms.

Ankyrin repeat (AR) proteins, first isolated in mammalian erythrocytes, are involved in the targeting, mechanical stabilization and orientation of membrane proteins to specialized compartments within the plasma membrane and endoplasmic reticulum (Mosavi et al., 2004). By extension, ANKRD13A may act as an adapter to recruit further proteins to the Tg PV. The ability of ANKRD13A to bind to Lys63- ubiquitinated substrates, as well as its role in endolysosomal trafficking, drew our attention to this protein as having potential relevance to the K63-ubiquitin dependent endolysosomal destruction of Tg type II Pru that we have observed in primary endothelial cells (Clough et al., 2016). We validated by ANKRD13A knock down, that the protein indeed controlled Tg in IFNγ-stimulated HUVEC and employing immunofluorescence microscopy we verified its localization to the Tg type II Pru and type III CEP PV.

ANKRD13A is an AR protein with four ubiquitin interaction motifs (UIMs) that specifically bind Lys63-linked ubiquitin and has been postulated to act as a ubiquitin-dependent regulator of epidermal growth factor receptor (EGFR) internalisation (Tanno et al., 2012). The tuning of ANKRD13A activities by ubiquitin modulation has been further highlighted by its interaction with the E3 ligases RNF11 and ITCH (Mattioni et al., 2020). Here, the authors showed that ANKRD13A acts as a scaffold allowing the transient formation of a complex between the E3 ligases ring finger protein 11 (RNF11) and Itchy E3 ubiquitin-protein ligase (ITCH) and EGFR, with the E3 ligases regulating the ubiquitination of ANKRD13A and thereby determining EGFR sorting for degradation or recycling. Tg has been reported to activate EGFR, albeit through an IFNγ-independent CD40 ligand-dependent mechanism, thus preventing autophagy-mediated killing of the parasite (Muniz-Feliciano et al., 2013). It is plausible to speculate that in type I infections and infected cells not stimulated with IFNγ, ANKRD13A is not ubiquitinated and can bind and mediate EGFR internalisation, thus enabling the parasite to evade host destruction via the acidification pathway. More recently, ANKRD13A has been implicated in the regulation of apoptotic cell death through its interaction with receptor interacting protein (RIP) 1 (Won et al., 2021). Here it binds ubiquitinated RIP1 preventing the recruitment of FADD and Caspase 8 to RIP1 preventing formation of cell death complex II. As mentioned above, ANKRD13A modulates ubiquitination of RNF11, an essential component of the A20 ubiquitin-editing protein complex, including ITCH and TAX1BP1 which can remove Lys63 ubiquitin from RIP1 (Jacque and Ley, 2009). TAX1BP1 has also been shown to be recruited to Tg type II Pru vacuoles in A549 lung epithelial cells and in MIO-M1 Müller glial cells ((Hernandez et al., 2022) and Clough et al. unpublished). Apoptotic cell death is employed in response to Tg infection of human macrophages as a mechanism of infection control and it is conceivable that cellular interactions involving ANKRD13A may skew the response to Tg infection in different cell types (Fisch et al., 2020).

In looking for ANKRD13A-associated proteins which may affect parasite survival, we noted an interaction with p97/VCP. ANKRD13A has been shown to facilitate the proteasomal degradation of caveolin-1 by forming a complex with p97/VCP, a hexameric AAA+ ATPase, and caveolin-1 (Burana et al., 2016). Its association with p97/VCP, a protein known for its ATP-dependent membrane segregase activity, led us to examine whether p97/VCP was also present at the PV and able to restrict *Toxoplasma*. We were able to verify that p97/VCP mediated IFNγ-dependent control on the Tg type II Pru and type III CEP and localised to the PV.

p97/VCP has many functions including the extraction of ubiquitinated proteins from membranes and serving as an interaction hub for >30 cofactors (Meyer and Weihl, 2014). Furthermore, p97/VCP is involved in modulating pathogen survival in both *Salmonella* and *Legionella*. In the former, a bacterial protein SptP regulates the activity of host p97/VCP by dephosphorylation, leading to increased intracellular replication of *Salmonella* (Humphreys et al., 2009). In the latter, p97/VCP was observed to promote bacterial replication by regulating the turnover of ubiquitinated proteins at the *Legionella*-containing vacuole (Dorer et al., 2006). p97/VCP may similarly modify the protein architecture at the PV of Tg by removing ubiquitinated pathogen virulence or protective proteins. The function of p97/VCP is governed by its interaction with specific cofactors that mediate its subcellular localisation as well as directing substrate binding and p97/VCPs function in cellular processes and pathways (Buchberger et al., 2015). Although the mammalian p97/VCP cofactors number ∼40, there are three major co-factors; p47, UFD1-NPL4 and UBXD1. Of these, UBXD1 has been linked to endolysosomal trafficking (Ritz et al., 2011) and has been described as a cofactor for VCP in a range of co-operative roles including the endolysosomal damage response pathway (ELDR) (Papadopoulos et al., 2017), mitophagy (Bento et al., 2018), endoplasmic reticulum associated degradation (ERAD) (Nagahama et al., 2009), which links to our hypothesis of endolysosomal pathway for Tg PV acidification. Further to this, we were unable to localise p47 to the PV in IFNγ-stimulated HUVEC, whereas UBXD1 was recruited to p97/VCP-positive vacuoles as shown in Figure 2e.

To date few universal host-defence operators have been observed at the PV in human cells. Notably ubiquitin leads this vacuole-situated pathogen control, with p62 a prominent player in all cell types studied. The mechanistic role of p62 in different cell types, however, is unclear and may diverge between those cells where vacuole breakage and cell death predominate versus those restricting Tg within its PV. Similarly, in this study we show that p97/VCP is a host protein that not only targets PVs in different human cell types, but additionally functions in Tg control. Since we observe varying levels of Tg survival with the knock down of p97/VCP in different cells, this host protein may participate in Tg restriction through alternate pathways or at different stages of Tg and PV elimination.

Together our findings suggest that a complex balance of ubiquitination and protein remodeling may be occurring at the PV, involving localized ANKRD13A, p97/VCP and UBXD1 that could direct pathogen control towards different pathways dependent upon cell type.

## EXPERIMENTAL PROCEDURES

List of reagents used in the study is provided in Supplementary Table S1

### Cell and parasite culture

Cultures of A549 lung alveolar epithelial cells (from Max Gutierrez) and Human Foreskin Fibroblast (HFF) cells (ATCC #SCRC-1041) for maintenance of *Toxoplasma*, were grown in Gibco^TM^ DMEM with GlutaMAX (Thermo Fisher Scientific #12077549) supplemented with 10% heat inactivated FBS (Life Technologies) and cultured at 37°C in 5% CO_2_. Human Umbilical Vein Endothelial Cells (HUVEC, Promocell #C-12203) were cultured in Gibco^TM^ Medium 199 (Thermo Fisher Scientific #11554426) supplemented with 20% heat-inactivated FBS, Heparin (Merck) and endothelial cell growth supplement (ECGS, Merck #02-102). THP-1 monocyte cell line (TIB202, ATCC) was maintained in Gibco^TM^ RPMI with GlutaMAX (Thermo Fisher Scientific #61870036) with 10% heat-inactivated FBS. THP1s were differentiated to macrophages with 50ng/ml PMA (Merck #P1585) for 3 days and then rested for 2 days in PMA-free, complete medium.

*Toxoplasma gondii* stably expressing luciferase/EGFP or Td-tomato (Prugniaud type II, and CEP type III) were maintained *in vitro* by serial passage on monolayers of HFF cells, cultured in Gibco^TM^ DMEM with GlutaMAX (Thermo Fisher Scientific) supplemented with 1% FBS at 37°C in 5% CO_2_. All cell culture was performed without addition of antibiotics. Cell lines and parasites were routinely tested for mycoplasma contamination by PCR test.

### Cell treatments

Cells were stimulated for 18-24h in complete medium at 37˚C with addition of 50- 100U/ml human IFNγ (R&D Systems #285-IF-100).

The E1 inhibitor PYR41 (Tocris #2978) was added to cells prior to infection at a concentration of 50μM and incubated at 37°C for 2h. Before infecting cells with Tg, the cells were washed twice with warmed PBS, so that the inhibitor did not impact ubiquitination within the parasite.

### Toxoplasma gondii infection

*Toxoplasma* were prepared from freshly 3x syringe-lysed (1×25G then 2×27G) HFF cultures. The parasite suspension was then briefly centrifuged at 50 xg for 3 mins to pellet and remove cell debris before adding parasites to experimental cells at a multiplicity of infection (MOI) of 1-2:1. The cell cultures with added *Toxoplasma* were then centrifuged at 800 xg for 5 min to synchronise the infection, prior to culturing at 37°C, 5% CO_2_ for the required time.

### Stable isotope labeling with amino acids in cell culture (SILAC)

SILAC Mass Spectrometry was used to quantitate ubiquitination of host cell proteins in epithelial cells infected with Tg. A549 were cultured in SILAC heavy and light media containing 100mg/L L-proline (DMEM containing ‘heavy’ ^13^C-Lys and ^13^C-Arg, #K8R10 or ‘light’ ^12^C-Lys and ^12^C-Arg, #K0R0 and supplemented with 10% dialysed FBS), for six cell doublings to allow incorporation of ^13^C-amino acids [Ong and Mann, 2007]. Both ‘heavy’ and ‘light’ A549 were transferred to corresponding SILAC media containing 0.5% dialysed serum for 12h before stimulating with 100U/ml IFNγ for 18h. The ‘light’ A549 cells were then infected with syringe-lysed, type II *Toxoplasma* MOI=4 for 2.5h after which infected cultures were washed with 3 x 10ml PBS to remove free parasites. ‘Heavy’ uninfected cultures and ‘light’ infected cultures were resuspended using trypsin EDTA and mixed in a 1:1 ratio. The cell mixture was lysed with 9M urea in 20mM HEPES pH8.0, supplemented with 1000U benzonase/10ml urea, and sonicated on ice (3mm probe, 50% amplitude, 3 x 15s bursts). Chloroacetamide was added to 20mM to inhibit deubiquitination enzymes. Protein concentrations were determined by Bradford Dye method (BioRad) with ∼30mg protein extracted in total. A final concentration of 10mM dithiothreitol (DTT)(Sigma) was added to reduce the proteins and incubated at 37°C for 30mins. Next, the extracted proteins were alkylated using 20mM chloroacetamide (Sigma) and incubated 30 min at RT in the dark. The alkylation reaction was quenched by addition of 10mM DTT for 10 min RT. Initial digestion with 15μg Lys-C (Promega) was performed by incubation 2h RT. A ∼20μg aliquot was taken to evaluate digestion by SDS-PAGE. The lysate was then diluted with 100mM ammonium bicarbonate, 5% acetonitrile to a final urea concentration of <2M. Samples were then digested by addition of trypsin 1:100 enzyme to protein (w/w) and incubated 37°C overnight. The following day the sample was digested two more times with trypsin (1:100, w/w) for 4h each at 37°C.

### Enrichment of di-Gly peptides

Peptides containing the di-Gly remnant were enriched using K-ε-GlyGly affinity resin (Cell Signalling Technologies #5562) according to manufacturer’s protocol. Briefly, digests were reconstituted in 1.4 ml of immunoaffinity purification (IAP) buffer as supplied by the manufacturer. One aliquot (∼40-µl packed bead volume) was washed four times with PBS and mixed with the peptide sample. Incubation of sample and beads was performed with gentle rotation at 4 °C for 2 hours followed by a 30 sec 2000 ×g spin to pellet the beads. The antibody beads were washed twice with ice-cold IAP buffer followed by three washes with ice-cold water. DiGly peptides were eluted from the beads with the addition of 50 µl of 0.15 % TFA and allowed to stand at room temperature for 5 min. After a 30 sec 2000 ×g spin, the supernatant was carefully removed and lyophilised for further LC-MS/MS analysis.

### LC-MS/MS

For mass spectrometry (MS) analysis, peptides were resuspended in 0.1 % TFA and loaded on 50 cm Easy Spray PepMap column (75 µm inner diameter, 2 µm particle size, Thermo Fisher Scientific) equipped with an integrated electrospray emitter. Reverse phase chromatography was performed using the RSLC nano U3000 (Thermo Fisher Scientific) with a binary buffer system at a flow rate of 250 nl/min. Solvent A was 0.1 % formic acid, 5 % DMSO, and solvent B was 80 % acetonitrile, 0.1 % formic acid, 5 % DMSO. The diGly enriched samples were run on a linear gradient of solvent B (2- 40 %) in 90 min, total run time of 120 min including column conditioning. The nanoLC was coupled to a Q Exactive mass spectrometer using an EasySpray nano source (Thermo Fisher Scientific). The Q Exactive was operated in data-dependent mode acquiring HCD MS/MS scans (R=17,500) after an MS1 scan (R=70, 000) on the 10 most abundant ions using MS1 target of 1 × 10^6^ ions, and MS2 target of 5 × 10^4^ ions. The maximum ion injection time utilized for MS2 scans was 120 ms, the HCD normalized collision energy was set at 28, the dynamic exclusion was set at 10 s, and the peptide match and isotope exclusion functions were enabled.

### MS Data processing

Data processing was performed with MaxQuant software (version 1.3.0.5) as described previously (Cox et al., 2009). Parent ion and tandem mass spectra were searched against *Toxoplasma gondii* database. A list of 247 common laboratory contaminants provided by MaxQuant was also added to the database. For the search the enzyme specificity was set to trypsin with maximum of three missed cleavages for the diGly dataset and two missed cleavages for the rest of the data. The precursor mass tolerance was set to 20 ppm for the first search (used for mass re-calibration) and to 6 ppm for the main search. Carbamidomethylation of cysteines was specified as fixed modification, oxidized methionines and N-terminal protein acetylation were searched as variable modifications. Di-glycine-lysine was added to the list of variable modifications. The datasets were filtered on posterior error probability to achieve 1 % false discovery rate on protein, peptide and site level. The mass spectrometry proteomics data have been deposited to the ProteomeXchange Consortium via the PRIDE (Perez-Riverol et al., 2022) partner repository with the dataset identifier PXD042937.

### siRNA knock down

A combination of three siRNAs for ANKRD13A (ThermoFisher silencer select # s40093, s40094, s40095), two siRNAs for VCP (ThermoFisher silencer select # s14765, s14767) or three siRNAs for UBXD1 (ThermoFisher silencer select # s37291, s37292, s37293) and siRNA non-targeting control (AM4635) were used to transfect cells. HUVEC were nucleofected (AMAXA) with siRNAs (100pmol) using HUVEC transfection reagent (VPB-1002, Lonza). Cells were used 48h after transfection.

### Protein lysates, SDS-PAGE and Immunoblot

Cells were washed 3 x in PBS 4°C, before lysis in 1%Triton-X100 in 25mM Tris HCl, pH 7.4, 5mM MgCl_2_, 150mM NaCl plus 1x complete protease inhibitor cocktail III, without EDTA (Calbiochem). Lysates were put on a rotator at 4°C for 1h, prior to centrifugation 18,000rpm 4°C. Protein supernatant concentrations were determined by Bradford Dye Assay (BioRad) and 10μg/lane separated by SDS-PAGE. Gels were dry blotted (iBlot, ThermoFisher) onto nitrocellulose membranes and probed with specified primary antibody diluted in PBS containing 5% BSA, 4°C overnight. Blots were washed 3 x 10 min in Tris-buffered saline plus 0.01% Tween 20 (TBS/Tween) before probing with relevant secondary antibody diluted in TBS/Tween. After washing 3 x 10min in TBS/Tween, blots were rinsed in TBS and developed with Immobilon Western Chemiluminescence HRP substrate (Millipore) and imaged on Imager A680 (Amersham).

### High Content Imaging and analysis

HUVEC were plated at 10K cells/well in 96 well plates (Imaging plates; Falcon black-walled #353219) and were later IFNγ-stimulated after cells had attached. After 18h, cells were infected with type II *Toxoplasma-*GFP at MOI=1-2 for 6h. Plates were washed in PBS to remove uninvaded parasites and fixed in 4% PFA 15 min. The cells were permeabilised with PB (0.2% BSA, 0.02% Saponin in PBS) for 30 mins before staining with Hoechst 33342 5ug/ml and Cell Mask Red (ThermoFisher #H32712). After washing twice with PBS the stained plates were imaged on Zeiss Cell Discoverer 7 using 20x objective and 0.5x magnification. Images were analysed using Zen Blue software and infection analysis performed by Host Response to Microbe Analysis pipeline (HRMAn) to obtain values for percentage infection and vacuole/cell ratio. For recruitment analysis, cells were plated and infected for 2h before fixation as above and were then permeabiised and stained with antibody according to the immunofluorescence protocol below. Recruitment was counted manually (>200 vacuoles per condition), since the HRMAN analysis pipeline had not been trained for these cells and stains.

### Immunofluorescence Microscopy

HUVEC were plated on coverslips (12mm, LJ1.5, ThermoFisher) in a 24-well plate and cultured, IFNγ–stimulated and infected with *Toxoplasma* as described above. All following steps were carried out at RT. The cells were washed with PBS and fixed with 3% paraformaldehyde in PBS for 15-20min. The fix was aspirated and Perm-Quench solution (50mM NH_4_Cl, 0.2% w/v saponin, Sigma 47036 in PBS) added as a wash and replaced with fresh Perm-Quench and incubated for 10-15min. The Perm-Quench solution was replaced with PGAS (0.2%w/v fish gelatin, Sigma G-7765, 0.02% w/v saponin, 0.02% w/v NaN_3_ in PBS). Cells were incubated for at least 5 min and fixed cells could be stored in PGAS at 4°C for several days. Antibody incubations were carried out in a humid box, inverting coverslips onto 50LJl drops of primary antibody, diluted in PGAS, and incubating for 1h at RT (except anti-LC3B: overnight incubation 1:100 at RT). Coverslips were washed in 3 x 1ml volumes of PGAS before incubating for a further 1h with second antibody, diluted in PGAS, at RT in the dark. Washes of 3 x 1ml PGAS and 2 x 1ml PBS followed by 1ml PBS containing 1μg/ml Hoechst 33342 (Life Technologies) and finally 2 washes in dH_2_O prior to mounting on glass slides with Mowiol 4-88 (Polysciences Inc.). Mounting medium was allowed to harden overnight. Slides were viewed on an SP5-invert Confocal microscope using x100 objective and analysed using LAS-AF software or by Structured Illumination Microscopy (SIM) on Zeiss Elyra 7 using x63 objective and Zen Black imaging software. Composite images were assembled using FiJi software.

### Live microscopy

HUVEC were transfected by nucleofection using Neon according to manufacturer’s protocol with pEGFP-p97, which was a gift from Hemmo Meyer (Addgene plasmid #85670; http://n2t.net/addgene:85670;RRID:Addgene_85670). After 24h, cells were stimulated with IFNγ and infected the following day with Tg type II Pru-td-Tomato. Imaging was carried out in an environmentally controlled chamber, 37°C, 5% CO_2_ using a Nikon LTTL widefield microscope. Data was collected and analysed using Micro-Manager and FiJi.

### Data handling, statistical measurements and evaluation

Numerical data was plotted using Graph Pad Prism and presented with error bars as standard deviation. Significance of results was determined by 2-way ANOVA or 2 tailed Student t-test.

---

**Table 1.**
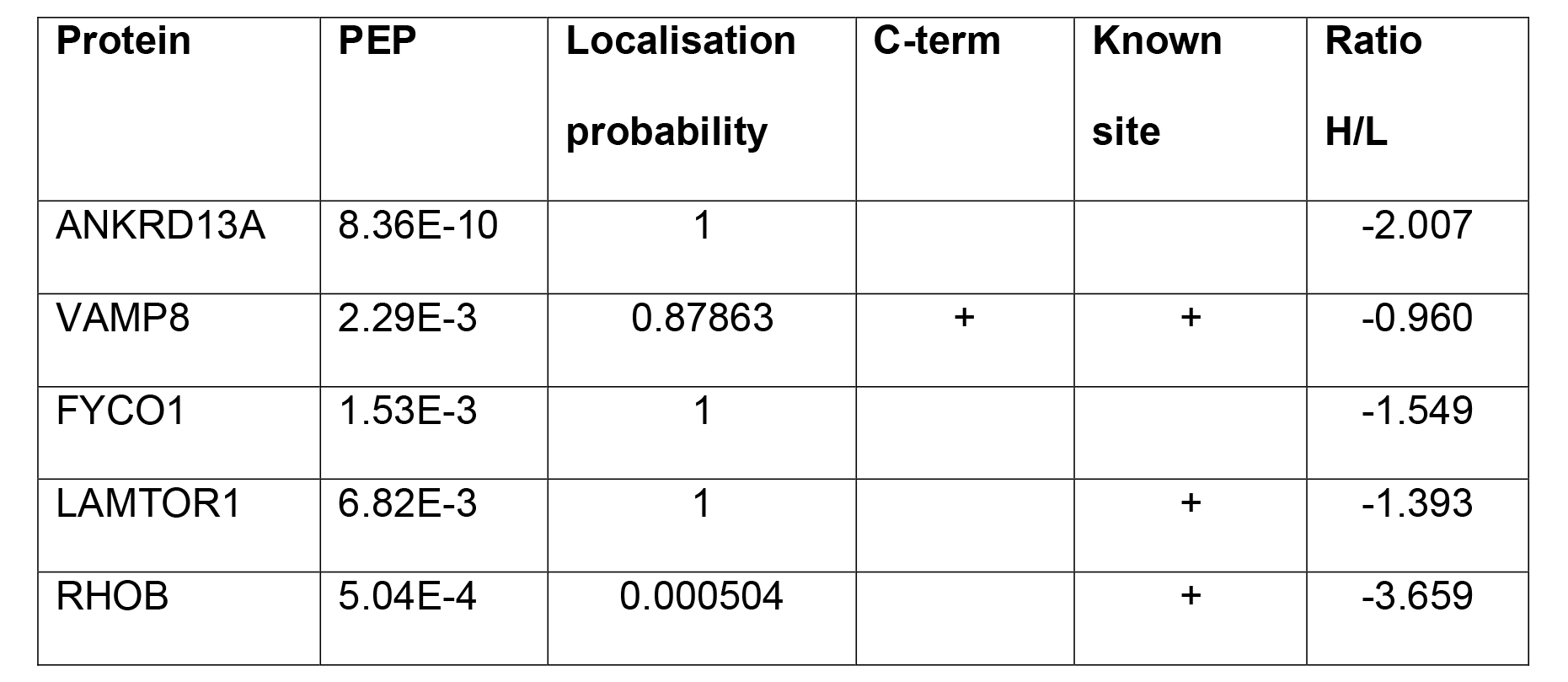
The identification, by LC-MSLS, of potential ubiquitin substrates listed with their respective peptide data. Posterior error probability (PEP) gives the probability that observed peptide spectrum matches is incorrect.

## Supplementary Figure Legends

**FIGURE S1. Mass Spectrometry identifies ANKRD13A as a ubiquitinated protein during *Toxoplasma* (Tg) infection.**

**a)** Representative structured illumination microscopy (SIM) image showing that the vacuoles of Tg type II Pru (white) in epithelial (A549) cells is ubiquitinated (red) in interferon gamma (IFNγ)-stimulated cells, nuclei (blue). Scale bar = 10μm.

**b)** A scheme for the stable isotope labelling by amino acids in culture (SILAC) workflow to identify substrates of ubiquitin is represented.

**c)** Intensity plot showing heavy/light ratio, for ubiquitinated peptides, with proteins of interest highlighted in red.

**FIGURE S2. Control of gene knockdowns using siRNA interference.**

**a)** Immunoblots and quantitation showing protein levels in interferon gamma (IFNγ)- primed HUVEC treated with control or target-specific siRNA. Representative blots of 3-5.

**b)** Immunoblots and quantification of protein expression after control and siRNA knockdown of VCP/p97 in HFF, HeLa and THP1 cells. Representative blots of 3.

**FIGURE S3. Recruitment kinetics of p97/VCP to *Toxoplasma* (Tg) type II Pru vacuoles.**

Live microscopy showing dynamic recruitment of EGFP-p97/VCP (green) to Tg type II Pru vacuoles (red) from 2h-2h50’ p.i. in interferon gamma (IFNγ)-stimulated HUVEC. Two cells i) and ii) are shown.

**FIGURE S4. Vacuolar acidification of Tg type II Pru at 6h p.i.**

Representative stitched structured illumination microscopy (SIM) image showing LysoTracker (red)-stained acidification of Tg type II Pru (white) vacuoles 6h p.i. in IFNγ-stimulated HUVEC. Scale bar = 30μm.

## Acknowledgements

The authors would like to acknowledge all members of the Frickel lab (University of Birmingham) for helpful discussions. They thank Matt Renshaw and Donald Bell in Crick Advanced Light Microscopy, The Francis Crick Institute, and Alex Di Maio in Birmingham Advanced Light Microscopy, University of Birmingham for advice and support with microscopic imaging. Further thanks are extended to Helen Flynn and Mark Skehel in the Crick Proteomics Science Technology Platform, The Francis Crick Institute, for helpful discussion and access to proteomics data.

## Contributions

BC and EMF conceived the study. BC performed all imaging and functional experiments. DF analysed imaging data with HRMAn. VE performed mass spectrometry. TM analysed mass spectrometry data. AS provided essential experimental protocols and equipment. EMF and AS acquired and contributed funding and experimental oversight. BC, DF & EMF analysed the data. BC and EMF wrote the manuscript, with input from all authors.

