## Supplementary figures and images for "p97/VCP targets *Toxoplasma gondii* vacuoles for parasite restriction in interferon-stimulated human cells"

### Supplementary Figures

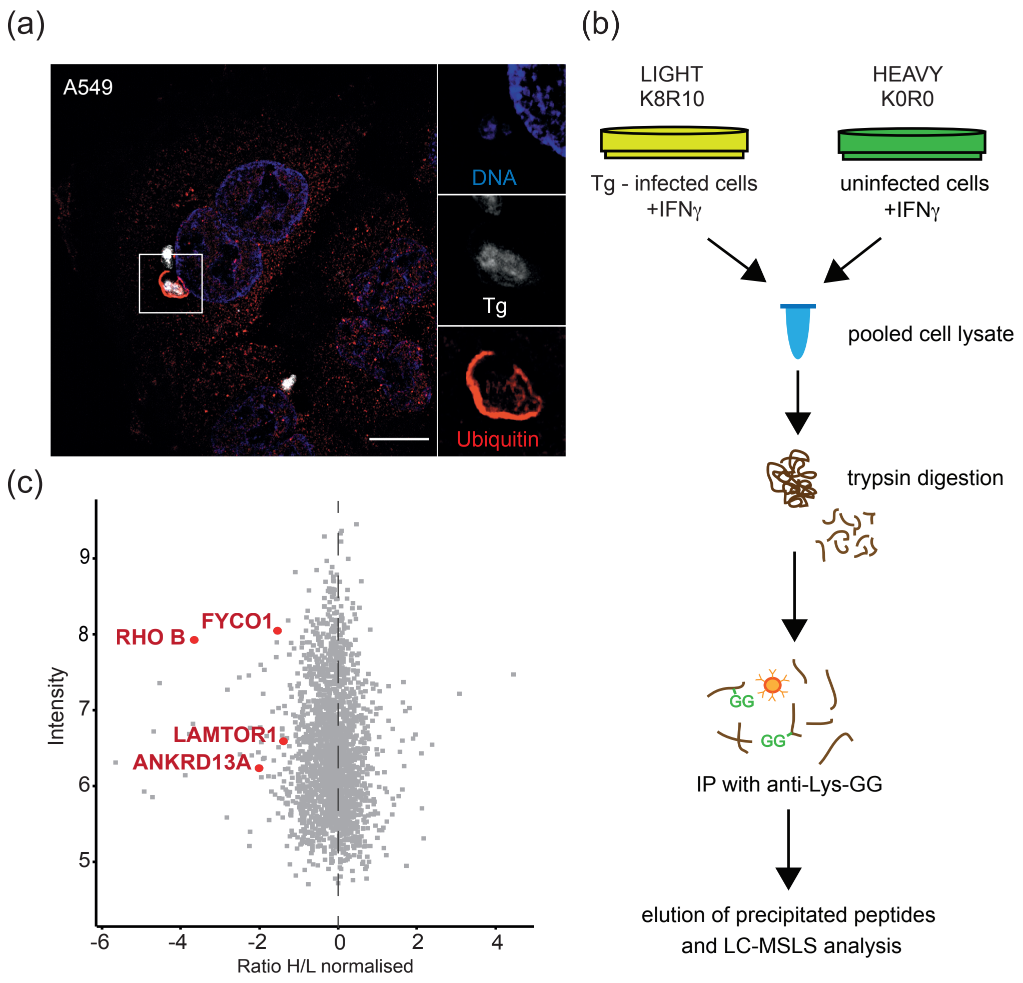

FIGURE S2

(a) HUVEC

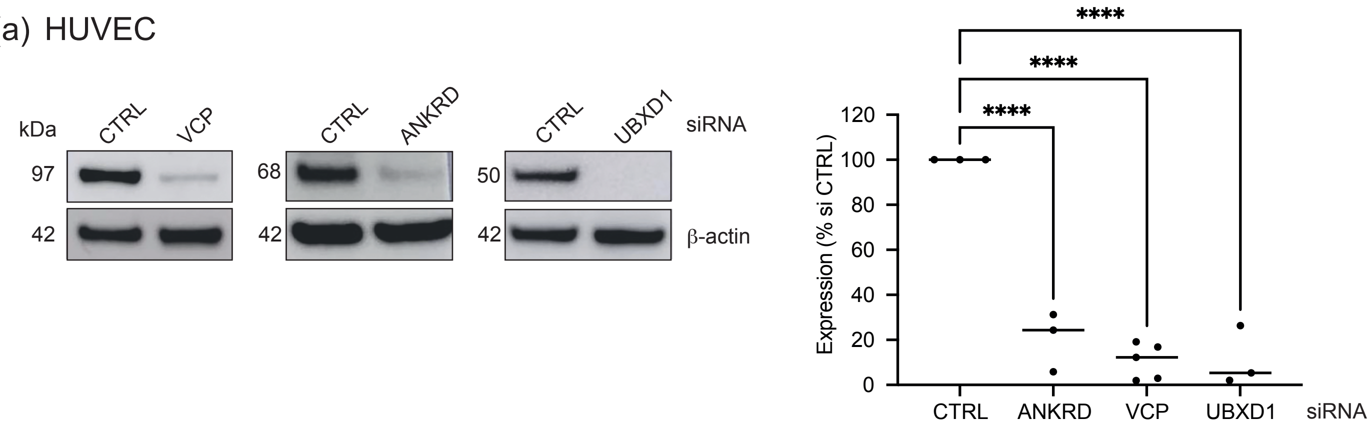


(b) HFF

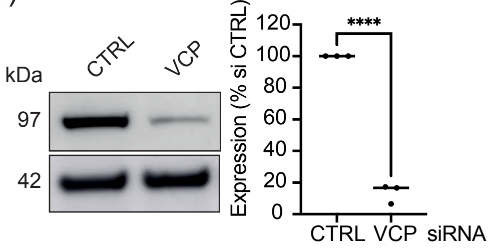


HeLa

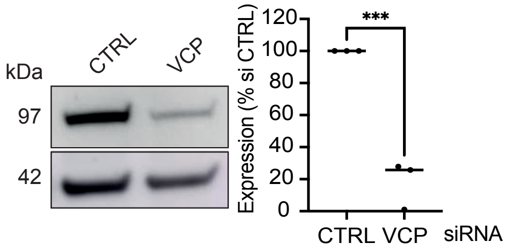


THP1

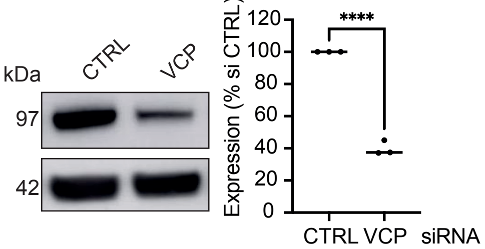

FIGURE S3

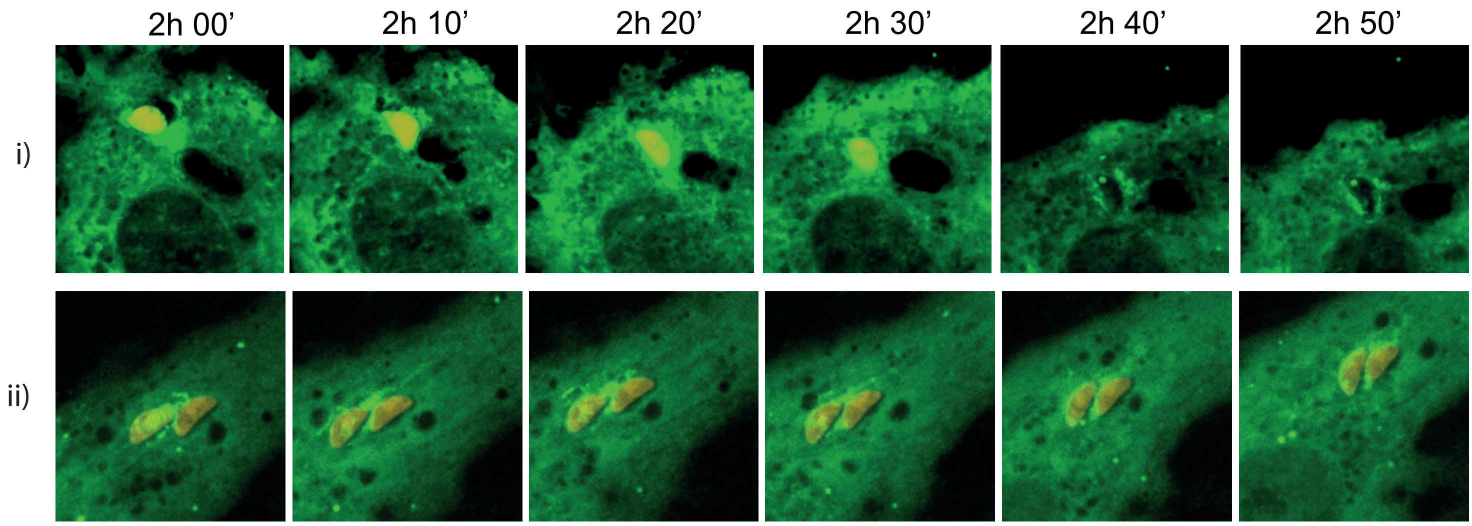


EGFP-p97/VCP 2-3h p.i. Tg type II Pru +IFNγ

FIGURE S4

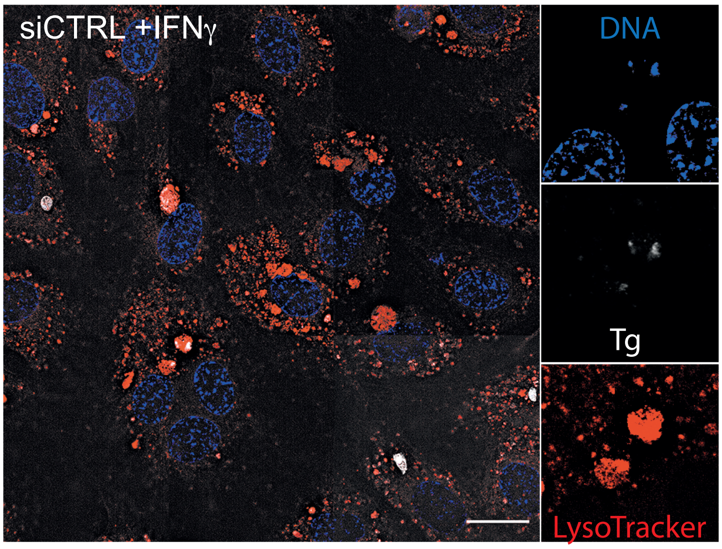
